# Spatiotemporal Variations in Growth Rate and Virulence Plasmid Copy Number during *Yersinia pseudotuberculosis* Infection

**DOI:** 10.1101/2020.11.04.369199

**Authors:** Stephan Schneiders, Tifaine Hechard, Tomas Edgren, Kemal Avican, Maria Fällman, Anna Fahlgren, Helen Wang

## Abstract

Pathogenic *Yersinia spp.* depend on the activity of a potent virulence plasmid-encoded *ysc*/*yop* type 3 secretion system (T3SS) to colonize hosts and cause disease. It was recently shown that *Y. pseudotuberculosis* up-regulates the virulence plasmid copy number (PCN) during infection and the resulting elevated gene dose of plasmid-encoded T3SS genes is essential for virulence. When and how this novel regulatory mechanism is deployed and regulates the replication of the virulence plasmid during infection is unknown. In the current study, we applied droplet digital PCR (ddPCR) to investigate the dynamics of *Y. pseudotuberculosis* virulence PCN variations and growth rates in infected mouse organs. We demonstrated that both PCN and growth varied in different tissues and over time throughout the course of infection, indicating that the bacteria adapted to discrete microenvironments during infection. The PCN was highest in Peyer’s Patches and caecum during the clonal invasive phase of the infection, while the fastest growth rates were found in the draining mesenteric lymph nodes. In deeper, systemic organs, the PCN was lower and more modest growth rates were recorded. Our study indicates that increased gene dosage of the plasmid-encoded T3SS genes is most important early in the infection during invasion of the host. The described ddPCR approach will greatly simplify analyses of PCN, growth dynamics, and bacterial loads in infected tissues, and will be readily applicable to other infection models.

**Importance:** Studying pathogenic bacteria proliferating inside infected hosts is challenging using traditional methods, especially the transit and reversible genetic events. The bacteria are effectively diluted by the overwhelming number of host cells present in infected tissues. Using an innovative droplet digital PCR (ddPCR) approach, we have determined the virulence plasmid copy number (PCN) variations and growth rates of *Yersinia* during the course of infection in a mouse model. Here, we show that both the virulence plasmid copy number and bacterial growth rates display spatiotemporal variations in mice during infection. We demonstrate that the peak-to-trough ratio can be used as a proxy for determining the growth rate of invasive bacterial pathogen during infection, and ddPCR as the method of choice for quantifying DNA in host-pathogen interaction context. This proof-of-concept ddPCR approach can be easily applied for any bacterial pathogens and any infection models, for analysis of PCN, growth dynamics and bacterial loads.

## Introduction

Bacterial pathogens have evolved various virulence strategies to successfully invade hosts and cause diseases. Pathogenic *Yersinia* deploys a potent plasmid-ncoded *ysc*/*yop* T3SS that inhibits phagocytic cells and suppresses the inflammatory response[1,2]. *Yersinia* virulence requires an increased gene dosage of plasmid-encoded T3SS genes through elevated PCN. This recently discovered regulation of an independently replicating plasmid is an essential virulence mechanism in *Yersinia*[3]. The actions of the T3SS and other virulence factors enable pathogenic *Yersinia* to evade the host immune response and proliferate in extracellular foci or microcolonies in lymphatic, liver, and spleen tissues[4–7].

While essential for virulence, deployment of the plasmid-encoded T3SS is metabolically expensive and pathogenic *Yersinia* species display calcium-dependent growth at 37°C due to the activity of the virulence plasmid-encoded T3SS[8]. At 37°C in the absence of Ca^2+^, pathogenic *Yersinia* switch on their T3SS and secrete massive amounts of bacterial toxins out of the cells[9,10]. Activation of T3SS secretion *in vitro* results in growth arrest and increased virulence PCN, and repression of T3SS (addition of excess Ca^2+^) restores bacterial growth. Thus, the activity of the T3SS is paradoxical and pathogenic *Yersinia* must have evolved mechanisms to trade-off essential virulence associated metabolic costs to successfully spread and proliferate during infection.

*Y. pseudotuberculosis* is a gastrointestinal pathogen that usually causes self-limiting gastroenteritis in humans[11,12], while high infection doses cause lethal systemic infection in mice[13,14]. Upon ingestion by mice, *Y. pseudotuberculosis* crosses the intestinal mucosal barrier, probably through microfold (M) cells in the intestinal epithelium, and proliferates in Peyer’s patches (PP) and the caecum[15,16]. Some bacteria drain to the mesenteric lymph nodes (MLN), followed by systemic spread to the liver and spleen[3]. The virulence plasmid-encoded T3SS is essential for the efficient systemic spread giving rise to the lethal infection[17], and T3SS-defective bacteria are still invasive but typically do not disseminate from the MLN[18].

Most bacteria bi-directionally replicate their circular chromosome from a single origin of replication (*oriC*). Therefore, within an actively replicating bacterial population, as the replication forks progress along the DNA molecule, the DNA dosage will be higher for genes closer to the *oriC* compared to genes closer to the replication terminus (*ter*) (Fig. 1A). For logarithmically growing bacteria, this difference in DNA dosage is evident in whole-genome sequencing (WGS) reads aligned to the circular chromosome, and a clear V-shaped coverage can be seen when the reads are aligned to the linearized chromosome (grey tracks in Fig. 1B, C). The growth rate of a bacterial population can be estimated by the ratio of the DNA coverage at the *oriC* divided by the coverage at the replication terminus[19,20]. This so-called peak-to-trough ratio (PTR) varies between 1 and 2 depending on the replicative status of the bacteria, and ratios of up to ~2.7 have been reported for fast-growing *Escherichia coli in vitro*[19].

**Fig. 1.**
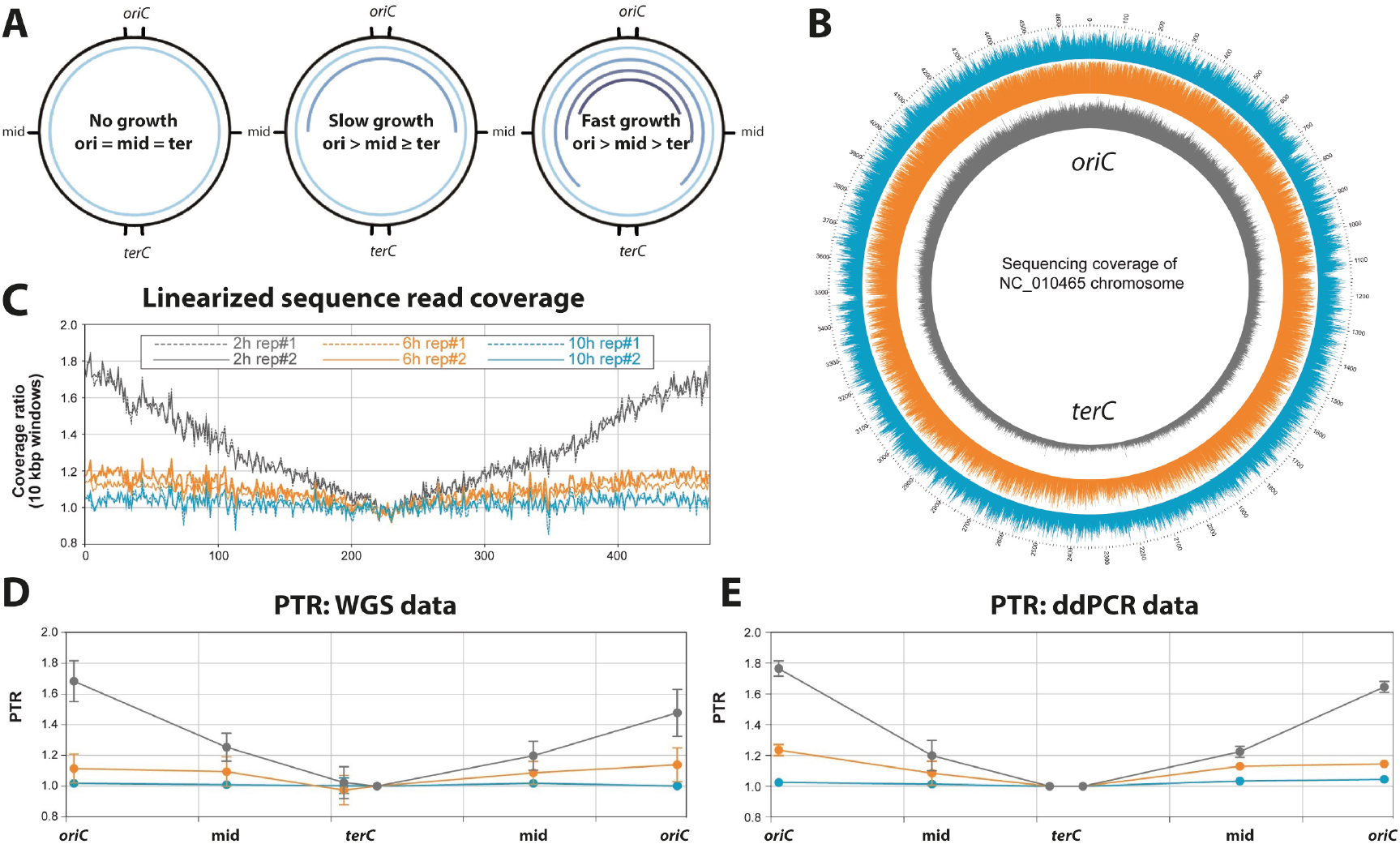
Droplet digital PCR (ddPCR) enables precise determination of peak-to-trough ratio (PTR) of bacterial populations. **A**, Bidirectional replication of circular chromosome leads to an enhanced DNA dosage of regions that have already been passed by the replication fork. Bacterial populations show different patterns of DNA dosage in chromosomal regions depending on their growth rate. Outer black lines indicate the positions of primer pairs used in this study. Inner blue circle lines indicate the DNA dosage of chromosomal regions. **B**, Sequence read coverage of *Y. pseudotuberculosis* DNA, isolated from a culture grown at 37°C with Ca^2+^ is plotted against a circular reference genome (*n* = 2). Grey, 2 h; orange, 6 h; blue, 10 h. **C**, Coverage of the same samples plotted against linearized reference chromosome. **D**, PTRs were calculated from the WGS data as the coverage depth at the position of the primers close to the origin divided by the coverage at the positions close to the terminus. **E**, PTRs of the same samples determined by ddPCR. **D.E**, Data represent the mean ± SD of two biological replicates.

In the current study, we optimized a cost-efficient, sensitive, and accurate droplet digital PCR (ddPCR)-based method for determining PCN and PTR in *Y. pseudotuberculosis*. ddPCR is a third-generation qPCR method that allows absolute quantification of the target DNA molecules present in a sample[21–23]. This method takes advantage of the random distribution of fragmented DNA in 20,000 water-oil emulsion droplets generated before the PCR reaction (Biorad). Each droplet is an independent PCR reaction, and determining the number of positive droplets enables accurate measurement of the number of target DNA molecules present in the sample. The PCN variations and growth rates of heterogeneous *Yersinia* communities proliferating at different infection sites have not been investigated previously. Therefore, after validating our ddPCR methodology, we investigated how the PCN and growth rates vary in *Y. pseudotuberculosis* populations in different tissues in a murine infection model.

## Results and discussion

### Development and validation of ddPCR for determination of PCN and PTR in *Y. pseudotuberculosis*

To validate our ddPCR method for establishing the PTR in *Y. pseudotuberculosis*, we determined the PTR of a *Y. pseudotuberculosis* YpIII:pIBX culture using both a PCR-free WGS approach (Fig. 1D) and ddPCR (Fig. 1E) with six different primer pairs targeting different regions of the chromosome (*oriC*, mid, and *ter* on either side of the *oriC*). The PTR of the *Y. pseudotuberculosis* culture ranged from a lowest value of ~1 for bacteria in stationary phase, to a highest value of ~1.8 at 2 h after inoculation. Critically, the PTR values calculated from ddPCR correlated with the PTR values determined by WGS of the same samples (Fig. 1D, E), validating the use of ddPCR for accurate PTR determination in bacterial cultures.

After verifying that the PTR of bacterial cultures could be determined by ddPCR, we next analysed how the PTR related to PCN and growth in *Y. pseudotuberculosis* cultures grown at 37°C under T3SS-repressed (+Ca^2+^) and T3SS-induced conditions (–Ca^2+^). Pathogenic *Yersinia* species display calcium-dependent growth at 37°C due to the activity of the virulence plasmid-encoded T3SS[8]. Activation of T3SS secretion *in vitro* results in growth arrest and increased PCN (black lines in Fig. 2A, B), and repression of T3SS (addition of excess Ca^2+^) restores bacterial growth and results in an increased PTR (Fig. S1). The T3SS-repressed culture (+Ca^2+^) displayed a PTR of 1.6–1.8 *oriC* equivalents per cell compared to the replication terminus during logarithmic growth (Fig. 2A, red dashed line). On the other hand, the T3SS-induced culture (−Ca^2+^) showed limited growth, with a PTR value of 1–1.2 throughout the experiment (Fig. 2A, black dashed line). In the T3SS-repressed culture, the PCN initially increased and then settled after 2 hours to around 2.5 virulence plasmid equivalents per cell, while in the T3SS-induced culture, the PCN increased to around 5 (Fig. 2B). We observed an inverse relationship between PTR and PCN *in vitro*; when the T3SS is active, *Yersinia* turn on PCN and growth slows down.

**Fig. 2.**
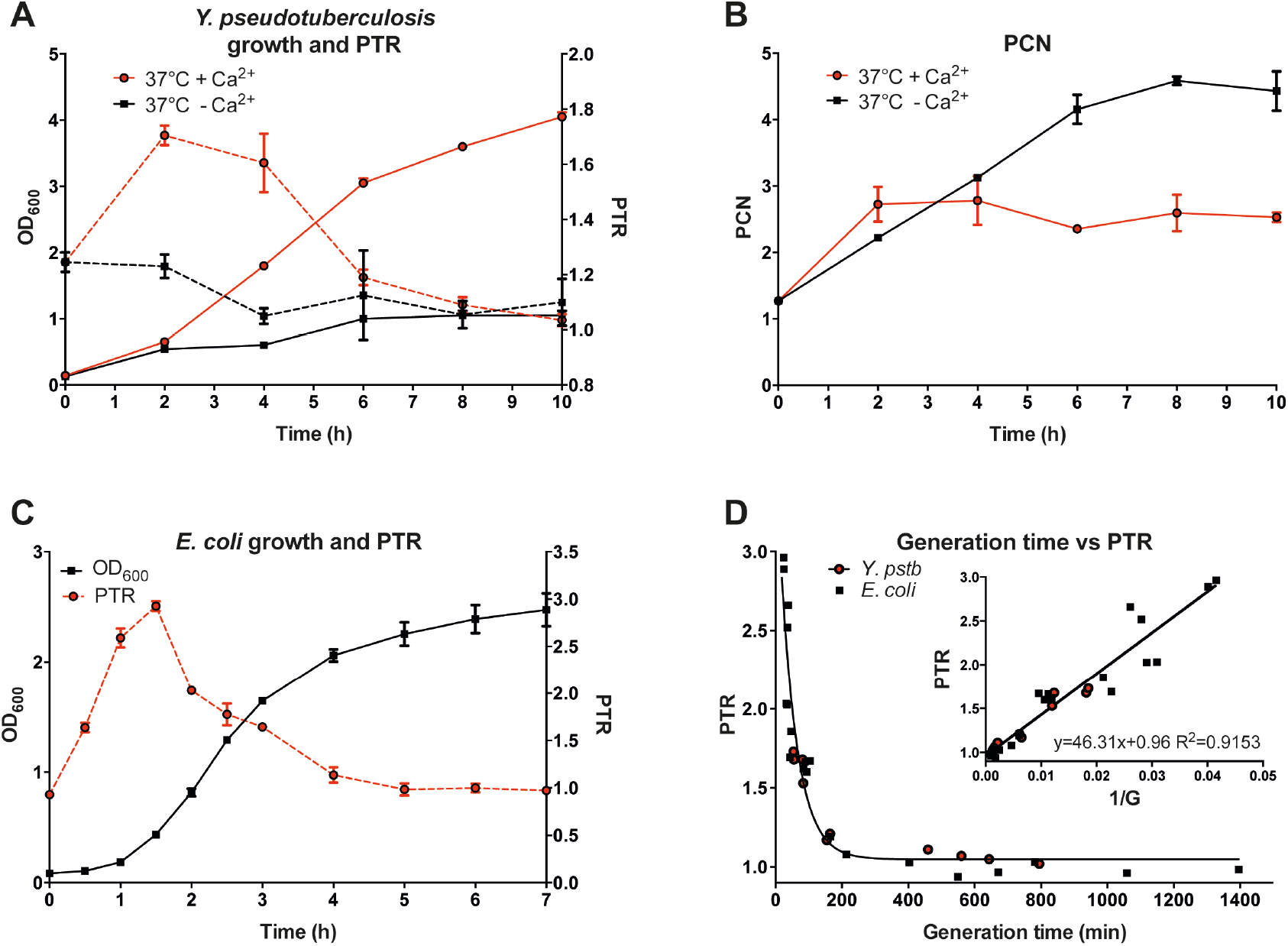
Droplet digital PCR (ddPCR)-based peak-to-trough ratio (PTR) reflects *in vitro* growth rates of *Y. pseudotuberculosis* and *E. coli*. **A.B**, Type 3 secretion system (T3SS)-dependent up-regulation of plasmid copy number (PCN) results in a metabolic burden and inhibits *Y. pseudotuberculosis* growth. **A**, PTRs (dashed lines) calculated based on ddPCR reflect the growth rate (solid lines) based on optical density of *Y. pseudotuberculosis* cultures grown under T3SS-repressive (37°C +Ca^2+^; red) and T3SS-induced (37°C −Ca^2+^; black) conditions. **B**, PCN changes determined by ddPCR of the same cultures. **C**, Growth and PTR measurements of *E. coli* grown in LB at 37°C. **D**, PTR plotted against generation times in *Y. pseudotuberculosis* and *E. coli*. Data represent the mean ± SD of two biological replicates.

### PTR values reflect the growth rates of different *enterobacteriaceae* with different generation times

To further investigate the relationship between PTR and growth rates in different *enterobacteriaceae*, we determined the PTR and generation times of *Y. pseudotuberculosis* and *E. coli* cultures. When grown in LB at 37°C, *Y. pseudotuberculosis* and *E. coli* have different growth rates with doubling times of around 60 and 20 minutes, respectively. We found that the calculated fastest generation times were 54 and 24.5 minutes (Fig. 2A, C) during early log phase for *Y. pseudotuberculosis* and *E. coli* (MG1655[24]) cultures, respectively. Independently of growth rate, the plot of PTR values against calculated generation times fit an exponential one-phase decay curve (Fig. 2D). Inverting the generation time, yielded a linear model (Fig. 2D insert). Thus, the relationship between PTR and growth rate was similar in both species, indicating that PTR can be used as a proxy for determining the growth rate of entero-bacterial populations.

### ddPCR is a robust method for determination of PCN and PTR in complex samples with interfering DNA

These initial experiments were performed using DNA isolated from laboratory grown bacterial cultures that lacked interfering host DNA. Using traditional WGS techniques, it is challenging to determine accurate PTR and PCN values for invasive bacteria in infected tissues due to the low amounts of bacterial DNA compared to host DNA present in the sample. Each diploid eukaryotic cell contains approximately 1000 times more genomic content compared to a bacterial cell—effectively diluting out the bacterial reads in WGS data. To further validate our ddPCR method, we tested the dynamic range of the assay in both the presence and absence of murine DNA (Fig. 3A–E; Fig. S2A,B). The assay was robust with respect to foreign DNA, and could accurately detect the target bacterial DNA in the presence of at least 800,000-fold excess of murine DNA (32ng/ul mouse DNA in Fig. 3E), using both the virulence plasmid primer pairs and chromosomal primer pairs. Externally added murine DNA in the sample did not affect the dose dependency of the assay, or reduce the absolute detection of target DNA (Fig. 3A–E). With our ddPCR method, increasing total dsDNA present in the sample was associated with increased background fluorescence, making it more difficult to differentiate between negative and positive droplets when using high total DNA concentrations (Fig. S2C). We were able to differentiate between positive and negative droplets with up to 32ng/ul mouse DNA in the sample, corresponding to 640ng mouse DNA (Fig. 3E). These results showed that ddPCR is a sensitive and robust method, with a high dynamic range enabling precise determination of target DNA, even in complex samples with a high excess of interfering foreign DNA.

**Fig. 3.**
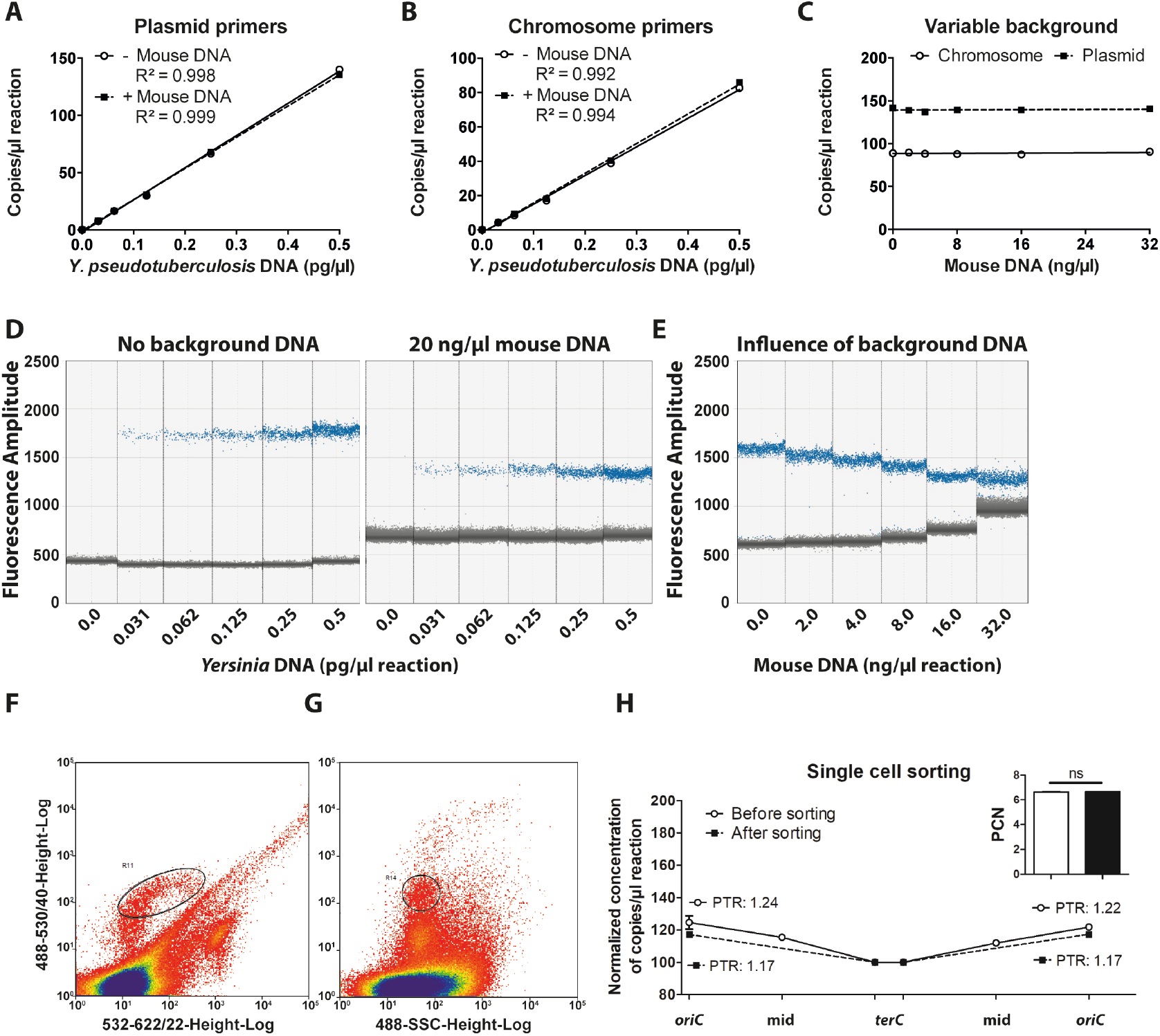
Droplet digital PCR (ddPCR) accurately determines the peak-to-trough ratio (PTR) and plasmid copy number (PCN) within a dynamic range of target and background DNA. **A-E.** Validation of ddPCR method using *Y. pseudotuberculosis* DNA isolated from a culture grown to stationary phase at 26°C. **A.B**, Plasmid and chromosome quantities in a dilution series of *Y. pseudotuberculosis* DNA, with and without 20 ng/μl mouse DNA as background. **C**, Plasmid and chromosome quantities from 0.5 pg/μl *Y. pseudotuberculosis* DNA with increasing levels of mouse DNA as background. **D.E**, Separation pattern of positive (blue) and negative (grey) droplets, **D**, with increasing concentrations of *Y. pseudotuberculosis* DNA, with and without 20 ng/μl mouse DNA as background, **E**, of 0.5 pg/μl *Y. pseudotuberculosis* DNA with increasing concentrations of mouse DNA as background. **F.G**, FACS sorting of homogenized and filtered Peyer’s patch (PP) sample taken 2 days post-infection, stained with SYBR green I and propidium iodide (PI). **F**, Gating of green fluorescence (SYBR, live) versus red fluorescence (PI, dead). **G**, Exclusion of highest SYBR green signals and large SSC (blue circle). **H**, Plotted coverage of normalized DNA quantity of the PP sample before sorting (white symbol; solid line) and after sorting (black symbol; dashed line) using ddPCR. Insert: PCN of the same sample before and after sorting. Data represent the mean ± SD of two technical replicates (ns = not significant, Student’s t-test).

### Determination of PCN and PTR in different tissues in a Murine infection model

To investigate the spatiotemporal variation of PCN and growth rates during infection, we infected mice with a virulent *Y. pseudotuberculosis* strain using an oral mouse infection model[3]. Mice were infected with 2 × 10^8^ colony forming units (cfu)/ml of the bioluminescent *Y. pseudotuberculosis* YpIII pIBX strain[16] via their drinking water. *Y. pseudotuberculosis* YpIII pIBX contains a mini-Tn5 *luxCDABE* inserted in the native pCD1 virulence plasmid, enabling visualization of bioluminescent bacteria in the infected organs using In Vivo Imaging System (IVIS) (Fig. S3). At different post-infection time-points, bioluminescent PP, caecums, MLN, spleens, and livers were collected from the infected mice. The larger organs (caecums, spleens, and livers) were further dissected to isolate the bioluminescent bacterial foci, in an effort to increase the amount of bacterial DNA compared to host DNA present in the samples.

We first conducted a control experiment designed to evaluate the accuracy of our ddPCR methodology in infected tissues. From infected PPs, bacterial cells were sorted from host cells using a MoFlo EQ flow cytometer (Beckman Coulter). The homogenized and filtered sample was stained with SYBR green I and propidium iodide (PI), enabling differentiation of PI-impermeable bacterial cells from dead eukaryotic cells and cell debris in the sample (Fig. 3F, G). A total of 24,141 bacterial cells were sorted, and we verified successful gating for *Yersinia* cells using microscopy and ddPCR quantification of estimated cell events. Importantly, we obtained similar PCN and PTR values from DNA isolated from the sorted cell population compared to DNA purified directly from the homogenates before sorting (Fig. 3H). This indicated that our ddPCR method can be used to accurately determine both the PCN and PTR in total DNA isolated from infected organs.

In PP at 1 day after infection, the PCN was ~6. This PCN level decreased with progressive infection, reaching ~2.5 in terminally ill mice that were euthanized at 6– 10 days after infection (Fig. 4A). The initial high PCN levels were similar to those previously reported using TruSeq WGS of PP at 2 days after infection[3]. In caecums, the PCN showed a similar but delayed trend, with a value of ~3 at 1 day post-infection, and a peak at ~6 at 2 days post-infection. Again, this PCN level decreased, and was ~3 in mice euthanized at 6–10 days post-infection. MLN samples showed a different trend, with the PCN remaining ~2.5 throughout infection, similar to the level observed in T3SS-repressed bacteria grown at 37°C. This is in line with previous studies showing that *Y. pseudotuberculosis* with a defective or absent *ysc*/*yop* T3SS can proliferate in MLN[18]. The deeper systemic organs also exhibited a relatively low PCN throughout the infection, with values of ~2 in spleens, and ~3 in livers. These findings are interesting since a functional T3SS is required for systemic spread of *Y. pseudotuberculosis* in mice. Moreover, systemic sites are colonized later in infection, and no bacteria were detected at 1 day post-infection and only two positive samples were identified at 2 days post-infection. Overall, we found that the PCN values at early infection sites (PP and caecum) were high in early infection, and decreased over time with progressive infection, i.e. after the bacteria had crossed the mucosa-associated lymphoid tissue and invaded deeper tissues (MLN, spleen, and liver). This suggests that high virulence PCN is probably most important during the clonal colonization phase compared to in later infection stages. Importantly, our results show that the PCN is not static during infection. The different PCN values in different tissues and over time indicate that the virulence PCN of *Yersinia* is regulated in response to environmental stimuli in different tissues.

**Fig. 4.**
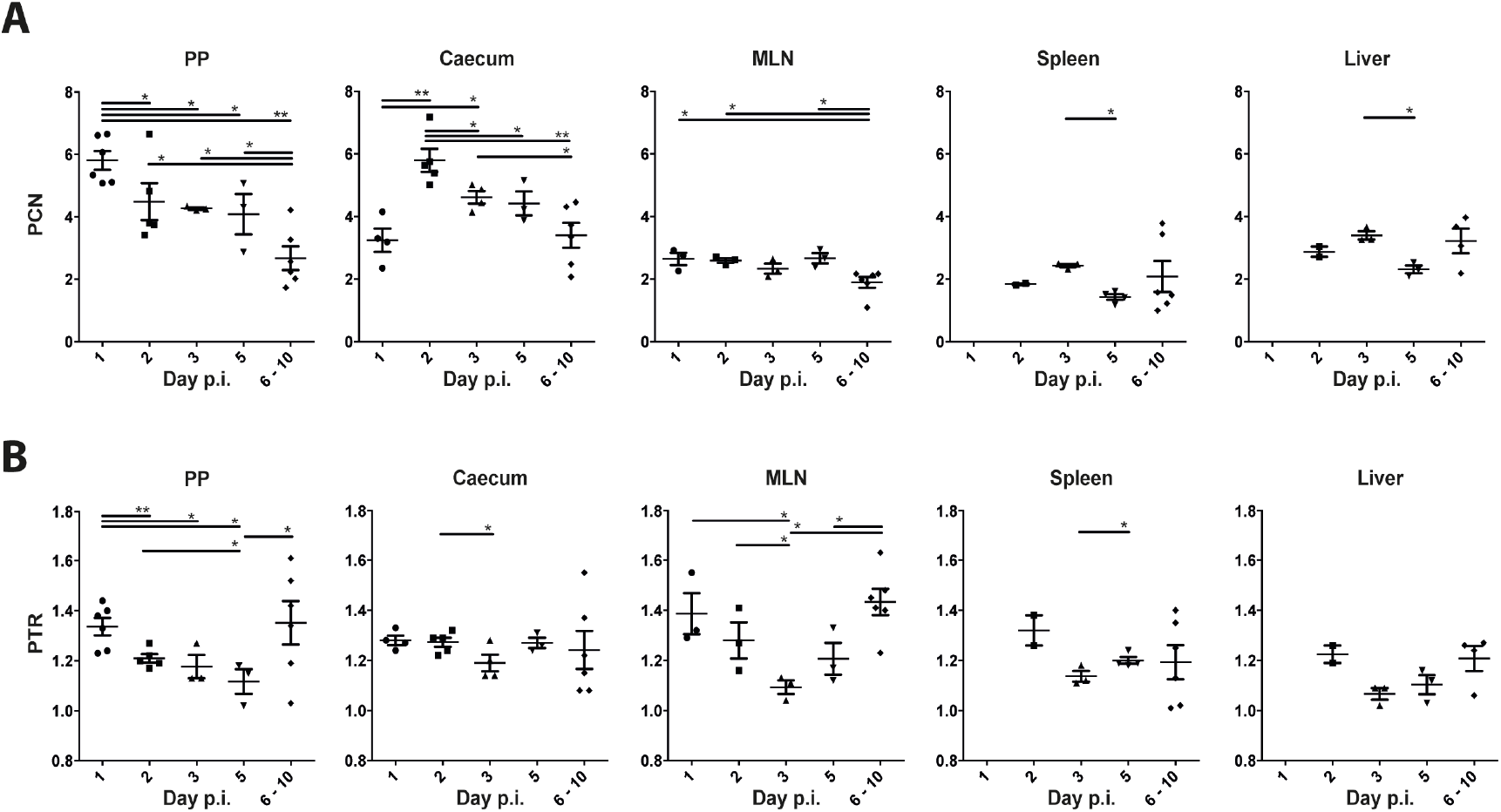
Spatiotemporal variations of PTR and PCN in mice during *Y. pseudotuberculosis* infection. **A**, Virulence PCN over a 10-day time period. PCN was calculated as the ratio between concentration of plasmid and chromosome DNA using ddPCR (*n* ≥ 2). **B**, PTR in different organs and at different time-points post-infection (p.i.). PTR was calculated as the ratio between DNA dosage of chromosomal regions located at *oriC* and *ter* (*n* ≥ 2) using ddPCR. Data represent the mean ± SEM of at least two biological replicates (\**P* ≤ 0.05; \*\**P* ≤ 0.01, Mann-Whitney U test).

We determined the PTR values of the infected samples using primer pairs proximal to the origin of replication and the replication terminus. In all examined samples, the PTR was between 1.1 and 1.4 (Fig. 4B), corresponding to generation times of ~600 and ~162 minutes, respectively. In comparison, we found PTR values of ~1.1 and ~1.8 for *Yersinia* grown *in vitro* under T3SS-repressed and T3SS-induced conditions, respectively (Fig. 2A). The recorded relatively low replicative rates during infection were expected, mirroring the struggle of invading bacteria to survive in the hostile host environment. We observed a tendency towards increased PTR values in organs from mice showing severe signs of infection and euthanized on days 6–10, compared to at 5 days post-infection. This indicates that the bacteria could proliferate faster in terminally ill mice after overcoming the initial host immune onslaught. We also observed a tendency for higher PTR values (~1.4) in PP and MLN at 1 day post-infection, which decreased to the minimum value at 5 days post-infection in PP, and 3 days post-infection in MLN. Overall, the growth rates as determined by PTR varies among different organs and over time, indicating that the bacterial populations confronted different microenvironments during the course of infection.

### *Y. pseudotuberculosis* have evolved mechanisms to trade-off virulence associated metabolic costs to maximize growth during infection

During infection, we saw no clear inverse relationship between PCN and PTR as seen in laboratory grown *Yersinia* cultures. Bacteria proliferating in PP at 1 day post-infection displayed high PCN values (~6) as well as high replicative rates (PTR of ~1.4). A similar decoupling of PCN and replication was also observed in the caecum at 2 days post-infection, where the PTR value was ~1.3 and the PCN values were also high. In the deeper tissues, the low PCN values recorded in MLN were associated with high replicative rates early in the infection (PTR of ~1.4 at 1 day post-infection), followed by slower growth at 3 days after infection. These data indicated that the T3SS-dependent growth restriction was not as evident during infection compared to in bacteria grown *in vitro*. This suggests that the invading pathogen must have evolved mechanisms to trade-off virulence-associated fitness costs to support proliferation in the hostile host environment.

In addition, the quantitative nature of ddPCR enables determination of the bacterial loads in the infected organs (Fig. S4). For our analyses, we isolated total DNA from whole PP and MLN, and DNA from dissected parts with bioluminescent foci from the larger organs (caecum, liver, and spleen). Early in the infection, we observed a delayed correlation between bacterial growth rates (PTR) and relative bacterial loads. For the first 3 days after infection, high PTR values were recorded, and the consecutive time-points showed increased bacterial loads. Later in the infection (day 5 and days 6–10), there was no clear relationship between PTR and bacterial loads. This discrepancy can most likely be explained by our analysis of only a portion of the dissected systemic organs in which bacterial growth would be expected at later time-points. The recorded bioluminescence emitted from whole mice in the IVIS images showed a similar trend, with a peak at 2 days after infection, followed by a decrease over time (Fig. S3).

### *Y. pseudotuberculosis* adapt to different micro-environments during infection

The presently observed differences in PCN and PTR among different organs and over time show that *Yersinia* colonization and infection is a dynamic process, during which bacteria must adapt to different environments. Enteric *Yersinia* species exhibit lymphoid tropism—colonizing PP, caecum, and MLN tissues[25]. Within these tissues, extracellular microcolonies are established in which the bacteria facing the host environment are exposed to phagocytes, while other bacteria are buried inside the microcolony. These different bacterial populations are clearly heterogeneous and display distinct gene expression profiles[5,25]. Our present findings support a model in which invading clonally expanding bacteria display high PCN and relatively high replicative rates early in the infection. Importantly, our data demonstrate *Yersinia* are replicating throughout the infection in all examined organs. As the bacteria proliferate and establish microcolonies, the relative importance of the T3SS decreases, which is reflected by the decreased PCN over time. The observed slower growth is most likely due to a combination of bacterial containment by the host immune system and nutritional limitation in the colony. The trend towards increased growth rates in terminally ill euthanized mice likely reflects the spread and proliferation of free bacteria as the infection progress. The dynamic changes of PCN in different infection sites has most likely evolved as a regulatory tactic to trade-off virulence costs during infection to maximize proliferation of *Yersinia* in the host. Regulation of plasmid-encoded functions by gene-dosage variations could be a general mechanism used by other bacteria in response to environmental cues.

**Fig. S1.**
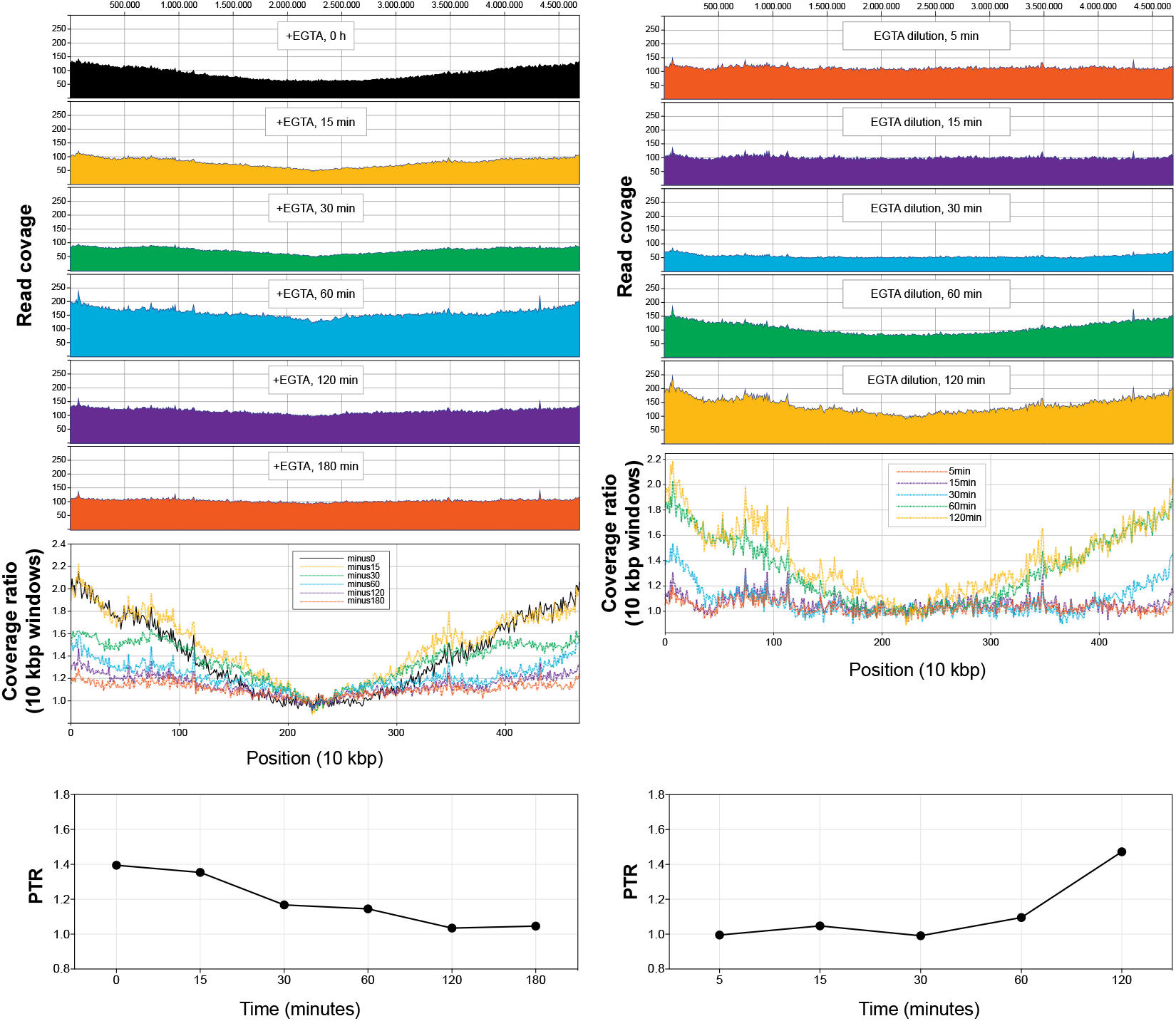
PTR of *Y. pseudotuberculosis* culture decreases under T3SS induction and can be reversed followed by repression of T3SS. Log-phase *Y. pseudotuberculosis* was shifted to T3SS-induced condition (37°C −Ca^2+^, “+EGTA”) for 3 hours, followed by addition of excess of Ca^2+^ (restoration of T3SS-suppression, “EGTA dilution”). Samples were taken at indicated time intervals and DNA was extracted and whole genome sequenced. Coverage depth was plotted against linearized reference chromosome. PTRs were calculated based on ddPCR methods describe above.

**Fig. S2.**
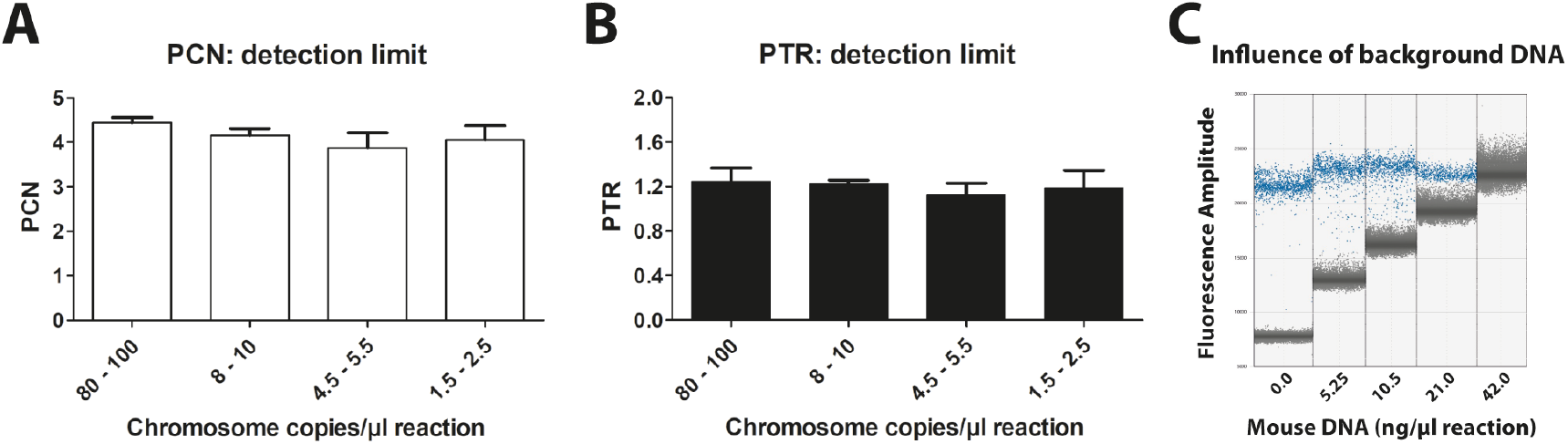
Determination of ddPCR detection limits. **A.B**, To determine the lower limit of target copies, dilution series of a Peyer’s patch sample taken 3 days post-infection (p.i.) were used to calculate the PTR and PCN. Data represent the mean ± SEM of two technical replicates. No significant differences were found by Student’s t-test. **C**, Separation pattern of 0.42 pg/μl *Y. pseudotuberculosis* DNA with an increasing concentration of mouse DNA as background.

**Fig. S3.**
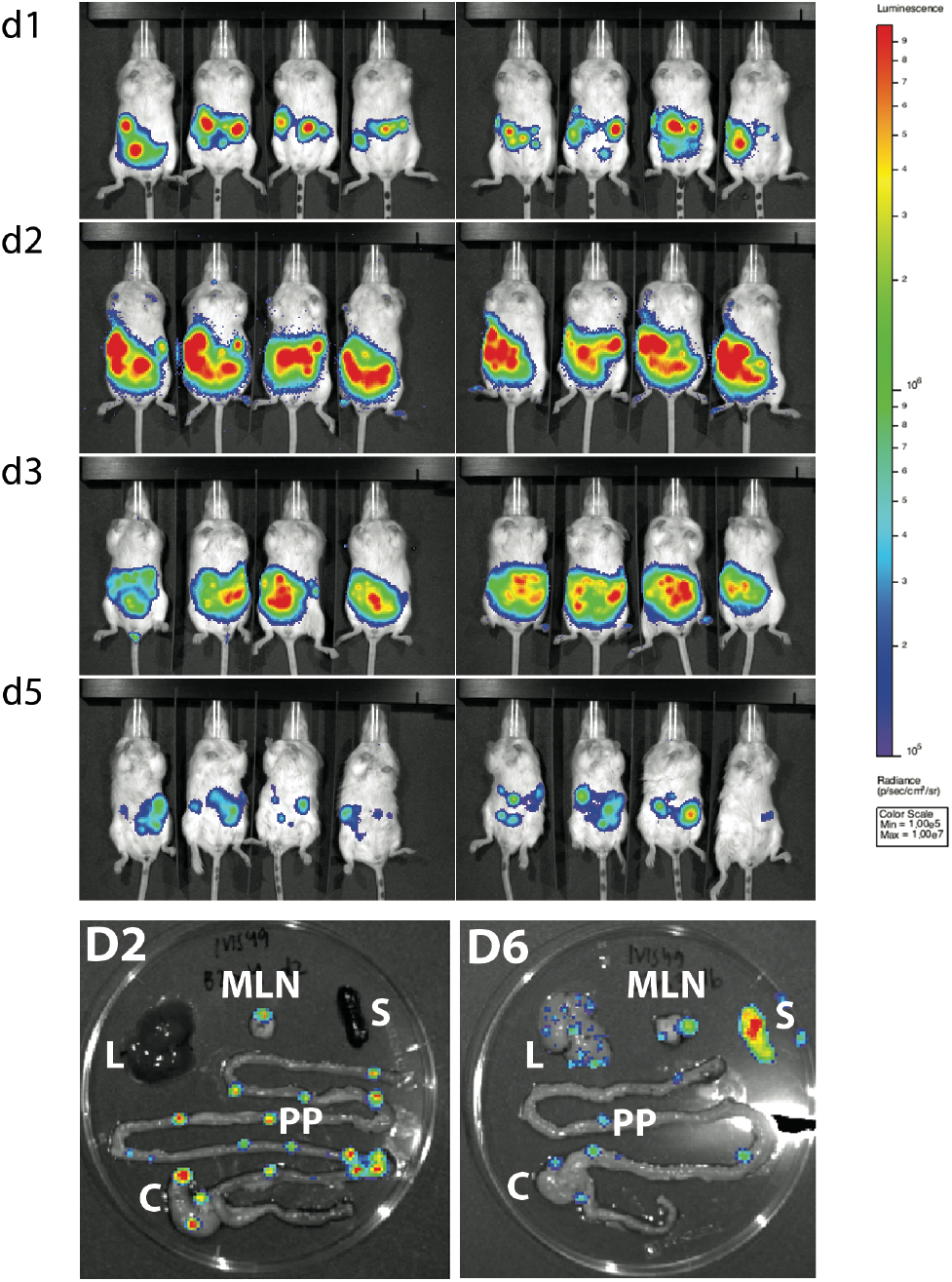
Monitoring of infections in mice and organ extraction using the In Vivo Imaging System (IVIS). *In vivo* images of mice infected with *Y. pseudotuberculosis*, and colours indicate levels of light emitted by bioluminescent bacteria. BALB/c mice (Charles River, 8-week-old, females) were orally infected with an infection dose of 2 × 10^8^ CFUs/ml. At indicated days after infection, mice were sacrificed and infected organs were dissected for DNA extraction and ddPCR analysis. Isolated organs: Peyer’s patches (PP), mesenteric lymph nodes (MLN), caecum (C), liver (L), and spleen (S).

**Fig. S4.**
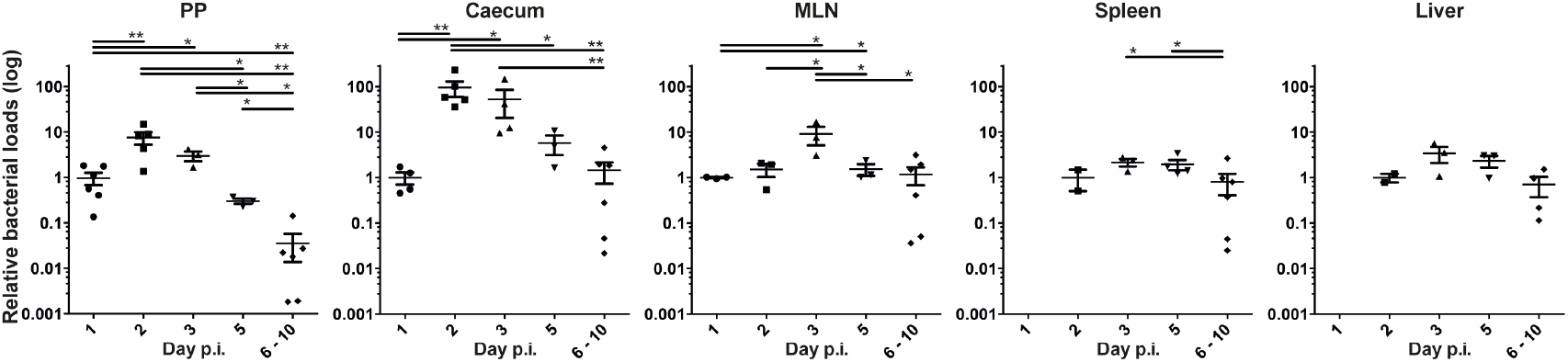
Determination of bacterial loads in infected mouse organs via ddPCR. The bacterial load per mg of infected organ tissue was calculated based on the copy number of chromosomal regions close to the terminus. Data were normalized against the value from first day post infection to show the relative changes of bacterial load during infection. Data represent the mean of at least two biological replicates (\**P* ≤ 0.05; \*\**P* ≤ 0.01, Mann-Whitney U test).

**Table S1.**
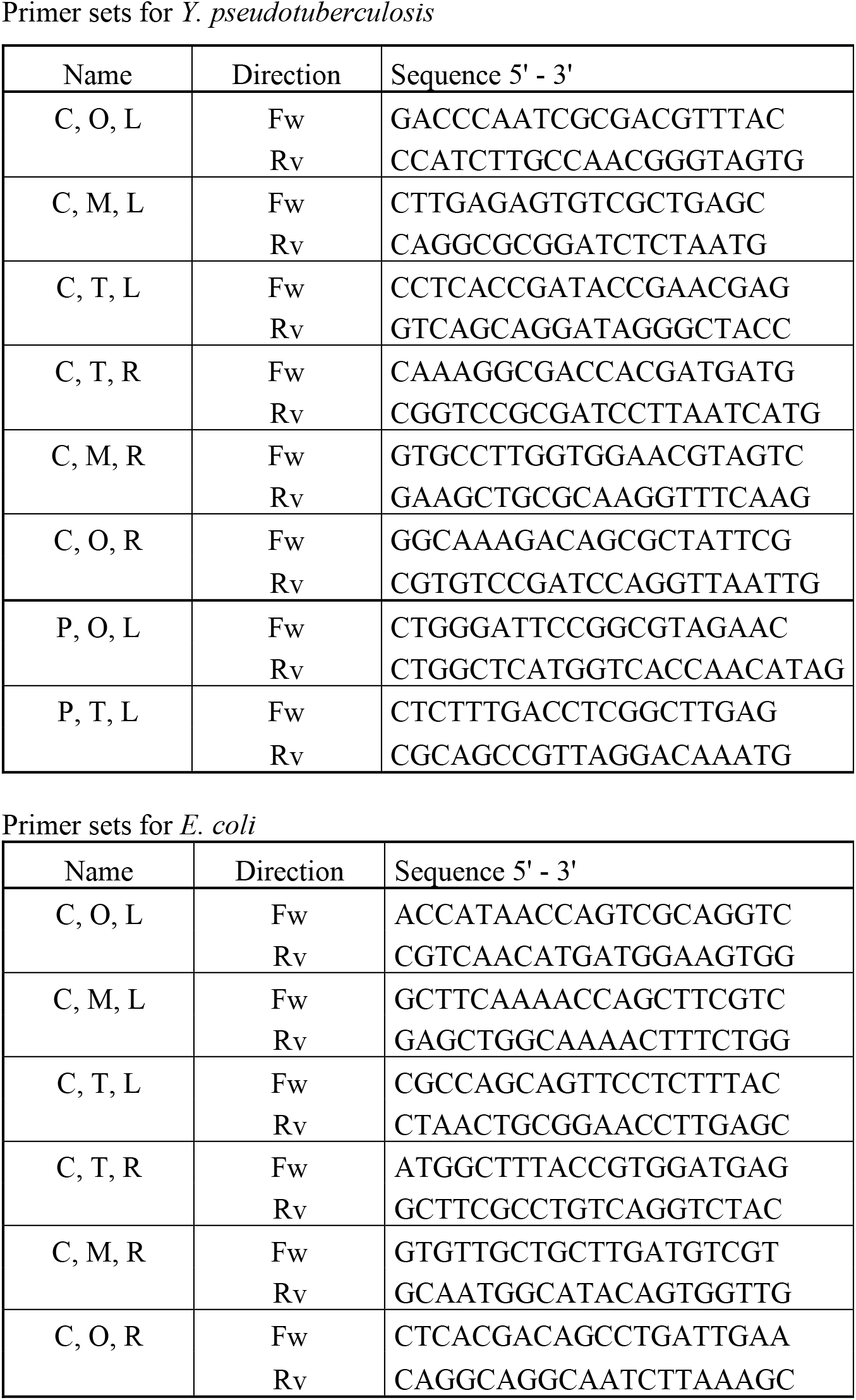
List of primers used in this study. (C = Chromosome, P = Plasmid, O = Origin, M = Middle, T = Terminus, L = Left, R = Right) Primer sets for *Y. pseudotuberculosis*

## Materials and Methods

### Bacterial Strains and growth conditions

*Yersinia pseudotuberculosis* YpIII/pIBX^*16*^ and *Escherichia coli* MG1655^*24*^ were routinely grown in Luria-Bertani (LB) broth or agar at 26°C for *Y. pseudotuberculosis* and 37°C for *E. coli*. Routine *Y. pseudotuberculosis* cultures were supplemented with 50 μg/ml Kanamycin (Km) for maintenance of the pIBX virulence plasmid. For T3SS induction experiments, overnight *Y. pseudotuberculosis* cultures were inoculated in fresh LB (1:20) supplemented with 5 mM CaCl2 (T3SS repressed conditions, +Ca^2+^) or 20 mM MgCl_2_ and 5 mM EGTA (T3SS induced conditions, - Ca^2+^) followed by 1 hour incubation at 26°C before shifting the cultures to 37°C. All growth analyses were carried out in duplicates and the bacterial densities were determined by measuring the optical density (OD) at 600 nm.

### DNA purification

Tissue samples were homogenized 2 × 55 sec in 1 ml sterile phosphate buffer saline (PBS) using a gentleMACS™ Octo Dissociator (Miltenyi Biotec). Total DNA from bacterial cells and infected tissues was purified using a GeneJet Genomic DNA Purification Kit (Thermo Scientific) according to the manufacturer’s recommendations for gram-negative bacteria and mammalian tissues respectively. The final concentration of dsDNA was determined using a Qubit 3.0 fluorometer according to manufacturer’s recommendations (Thermo Scientific).

### ddPCR

ddPCR analyses were carried out with EVA-Green using the Bio-Rad ddPCR QX200 system according to manufacturer’s recommendations. Briefly, 12.5 μl 2 x EVA-Green master mix (Bio-Rad), 1 μl HindIII-HF (NEB), 10.5 μl template DNA and 1 μl respective primer pairs (200 nM final concentration, supplementary table 1) were mixed and distributed into a 96-well qPCR plate (Bio-Rad). The plate was incubated 10 min at room temperature to digest the target DNA before droplets were generated in an Automated Droplet Generator (Bio-Rad). PCR was performed with a hot-start/enzyme activation at 95°C for 5 minutes, denaturation at 94°C for 30 seconds and amplification at 58°C for 1 minute over 40 cycles followed by signal stabilization at 4°C for 10 minutes and 90°C for 5 minutes. For all steps, a ramp rate of 2°C/s was used. Subsequently, the droplets were analyzed in the QX200 droplet reader (Bio-Rad). The data were analyzed with QuantaSoft™ Analysis Pro 1.0.569 (Bio-Rad).

### Whole genome sequencing and data analysis

Purified total DNA was sequenced using the Illumina TruSeq DNA PCR-free protocol by Novogene. Data analysis and Tracks generation were performed using the CLC Genomic Workbench 11 (CLC bio, Qiagen). Sequences were aligned against *Y. pseudotuberculosis* sequences (NC_010465 for the YPIII chromosome and an in house pIB1 sequence for the plasmid) using a 10 kb window. The PCN and PTR were calculated by dividing the average coverage of plasmid DNA with the average coverage of chromosomal DNA and the average coverage at *oriC* with the average coverage at *ter* of the mapped reads respectively. The sequence data used in this study have been deposited in the European Nucleotide Archive (ENA) with accession number PRJEB38239.

### Mouse infection and bioluminescent imaging

Eight-week-old female BALB/c mice (Charles river) were allowed to acclimate to the new environment for one week before the experiments. Mice were deprived of food and water for 16 hours prior to oral infection with bioluminescent wild-type YpIII/pIBX. Bacteria grown overnight at 26°C were harvested and re-suspended at a concentration of 2 × 10^8^ CFUs/ml in sterile tap-water supplemented with 150 mM NaCl. The bacterial suspensions were provided for 6 hours as drinking water. The infection dose was determined by viable counts and drinking volume. Mice were inspected frequently for signs of infection, and mice showing prominent clinical signs were euthanized promptly to prevent suffering. Mice were monitored for bioluminescent emission using IVIS Spectrum (Caliper LifeSciences). Mice were anesthetized using the XGI-8 gas anesthesia system (Caliper LifeSciences) prior to imaging with 2.5% IsoFluVet in oxygen (Orion Pharma Abbott Laboratories Ltd, Great Britain), and during imaging in 0.5% IsoFluVet. Images were acquired and analyzed using Living Image 4.5 (Caliper LifeSciences). To analyze bacterial localization within organs, mice were euthanized, the intestine, mesenteric lymph nodes, liver, and spleen removed, and the organs imaged by bioluminescent imaging. Bioluminescent regions of the larger organs (caecum, liver and spleens) were further dissected before freezing in liquid nitrogen.

Mice were housed in accordance with the Swedish National Board for Laboratory Animals guidelines. All animal experiments were approved by the Animal Ethics Committee of Umeå University (Dnr A65-15).

### FACS analysis and single cell sorting

A Peyer’s patch sample from 2 days after infection, was homogenized in 1 ml PBS for 2 × 55 sec on a gentleMACS™ Octo Dissociator (Miltenyi Biotec) using a C-Tube (Miltenyi Biotec), which allows dissociation of viable single cells from tissue samples. Single cells were isolated by filtering through a 30μm nylon mesh. Hereafter, an aliquot of the filtered sample was diluted 200 times and a live/dead staining was performed. For this purpose, samples were incubated for 10 minutes with SYBR Green I and propidium iodide (PI) to a final concentration of 1 μM and 2 μM, respectively.

The sorting was performed with a MoFlo Astrios EQ flow cytometer (Beckman Coulter) using the 488 nm laser and 530/40 nm filter (SYBR Green I; all cells), 532 nm laser and 622/22 nm filter (PI; dead cells) for excitation and emission, plus a 70 μm nozzle, sheath pressure of 60 psi and 0,1 μm filtered 1 x PBS as sheath fluid. Forward scatter was used as trigger channel. A tube sorting of single cells was performed with the most stringent settings (purify mode and 1-2 drop envelope) in order to obtain both a qualitative and quantitative measurement of the sorted sample. Cell sorting was performed at the Microbial Single Cell Genomics Facility at Science for Life Laboratory in Uppsala.

### Generation time calculations

The generation times of bacterial cultures were calculated from the 37°C growth curves presented in Fig 2 A and B for the T3SS repressed *Y. pseudotuberculosis* culture grown in the presence of Ca^2+^ (solid red line in Fig 2A) and *E. coli* (solid black line in Fig 2C) using the formula:

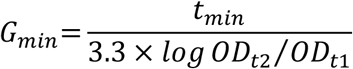

*t*_*min*_ is the time interval in minutes, *OD*_*t*1_ is the OD_600_ at the beginning of the time interval and *OD*_*t*2_ is the OD_600_ at the end of the time interval. Each calculated value from the biological duplicates was plotted against the corresponding PTR value at *OD*_*t*/2_. The resulting data fit a single exponential one-phase decay curve with a R^2^ value of 0.8872. A linear model was constructed by inverting the generation times (1/G) and both data sets fit a single line with the equation y=46.31x+0.96 with a R^2^ value of 0.9153.

### Quantification and Statistical Analysis

Statistical analysis details of each individual experiment are presented in the figure legends or relevant methods section. The statistical analyses were performed using GraphPad Prism 6.

## Acknowledgements

Sequence data used in this study have been deposited in the European Nucleotide Archive with accession number PRJEB38239 (available upon requests). This work is supported by Swedish Research Council grant 2018-02376 to HW, 2018-02855 to MF, Swedish Society for Medical Research Stora Anslag, Carl Tryggers Stiftelse för Naturevetenskapig Forskning, and Petrus and Augusta Hedlund Stiftelse grant to HW.

## Reference

1. Gemski P, Lazere JR, Casey T, Wohlhieter JA (1980) Presence of a virulence-associated plasmid in Yersinia pseudotuberculosis. Infect Immun 28: 1044–1047.

2. Portnoy DA, Wolf-Watz H, Bolin I, Beeder AB, Falkow S (1984) Characterization of common virulence plasmids in Yersinia species and their role in the expression of outer membrane proteins. Infect Immun 43: 108–114.

3. Wang H, Avican K, Fahlgren A, Erttmann SF, Nuss AM, et al. (2016) Increased plasmid copy number is essential for Yersinia T3SS function and virulence. Science 353: 492–495.

4. Oellerich MF, Jacobi CA, Freund S, Niedung K, Bach A, et al. (2007) Yersinia enterocolitica infection of mice reveals clonal invasion and abscess formation. Infect Immun 75: 3802–3811.

5. Davis KM, Mohammadi S, Isberg RR (2015) Community behavior and spatial regulation within a bacterial microcolony in deep tissue sites serves to protect against host attack. Cell Host Microbe 17: 21–31.

6. Logsdon LK, Mecsas J (2006) The proinflammatory response induced by wild-type Yersinia pseudotuberculosis infection inhibits survival of yop mutants in the gastrointestinal tract and Peyer’s patches. Infect Immun 74: 1516–1527.

7. Simonet M, Richard S, Berche P (1990) Electron microscopic evidence for in vivo extracellular localization of Yersinia pseudotuberculosis harboring the pYV plasmid. Infect Immun 58: 841–845.

8. Straley SC, Plano GV, Skrzypek E, Haddix PL, Fields KA (1993) Regulation by Ca2+ in the Yersinia low-Ca2+ response. Mol Microbiol 8: 1005–1010.

9. Michiels T, Wattiau P, Brasseur R, Ruysschaert JM, Cornelis G (1990) Secretion of Yop proteins by Yersiniae. Infect Immun 58: 2840–2849.

10. Bergman T, Erickson K, Galyov E, Persson C, Wolf-Watz H (1994) The lcrB (yscN/U) gene cluster of Yersinia pseudotuberculosis is involved in Yop secretion and shows high homology to the spa gene clusters of Shigella flexneri and Salmonella typhimurium. J Bacteriol 176: 2619–2626.

11. Hubbert WT, Petenyi CW, Glasgow LA, Uyeda CT, Creighton SA (1971) Yersinia pseudotuberculosis infection in the United States. Speticema, appendicitis, and mesenteric lymphadenitis. Am J Trop Med Hyg 20: 679–684.

12. Paff JR, Triplett DA, Saari TN (1976) Clinical and laboratory aspects of Yersinia pseudotuberculosis infections, with a report of two cases. Am J Clin Pathol 66: 101–110.

13. Heesemann J, Gaede K, Autenrieth IB (1993) Experimental Yersinia enterocolitica infection in rodents: a model for human yersiniosis. APMIS 101: 417–429.

14. Barnes PD, Bergman MA, Mecsas J, Isberg RR (2006) Yersinia pseudotuberculosis disseminates directly from a replicating bacterial pool in the intestine. J Exp Med 203: 1591–1601.

15. Clark MA, Hirst BH, Jepson MA (1998) M-cell surface beta1 integrin expression and invasin-mediated targeting of Yersinia pseudotuberculosis to mouse Peyer’s patch M cells. Infect Immun 66: 1237–1243.

16. Fahlgren A, Avican K, Westermark L, Nordfelth R, Fallman M (2014) Colonization of cecum is important for development of persistent infection by Yersinia pseudotuberculosis. Infect Immun 82: 3471–3482.

17. Galan JE, Wolf-Watz H (2006) Protein delivery into eukaryotic cells by type III secretion machines. Nature 444: 567–573.

18. Balada-Llasat JM, Mecsas J (2006) Yersinia has a tropism for B and T cell zones of lymph nodes that is independent of the type III secretion system. PLoS Pathog 2: e86.

19. Korem T, Zeevi D, Suez J, Weinberger A, Avnit-Sagi T, et al. (2015) Growth dynamics of gut microbiota in health and disease inferred from single metagenomic samples. Science 349: 1101–1106.

20. Brown CT, Olm MR, Thomas BC, Banfield JF (2016) Measurement of bacterial replication rates in microbial communities. Nat Biotechnol 34: 1256–1263.

21. Devonshire AS, Honeyborne I, Gutteridge A, Whale AS, Nixon G, et al. (2015) Highly reproducible absolute quantification of Mycobacterium tuberculosis complex by digital PCR. Anal Chem 87: 3706–3713.

22. Sze MA, Abbasi M, Hogg JC, Sin DD (2014) A comparison between droplet digital and quantitative PCR in the analysis of bacterial 16S load in lung tissue samples from control and COPD GOLD 2. PLoS One 9: e110351.

23. Huggett JF, Foy CA, Benes V, Emslie K, Garson JA, et al. (2013) The digital MIQE guidelines: Minimum Information for Publication of Quantitative Digital PCR Experiments. Clin Chem 59: 892–902.

24. Blattner FR, Plunkett G, 3rd, Bloch CA, Perna NT, Burland V, et al. (1997) The complete genome sequence of Escherichia coli K-12. Science 277: 1453–1462.

25. Davis KM (2018) All Yersinia Are Not Created Equal: Phenotypic Adaptation to Distinct Niches Within Mammalian Tissues. Front Cell Infect Microbiol 8: 261.

